# Deconvolution of the epigenetic age discloses distinct inter-personal variability in epigenetic aging patterns

**DOI:** 10.1101/2021.06.20.449142

**Authors:** Tamar Shahal, Elad Segev, Thomas Konstantinovsky, Yonit Marcus, Gabi Shefer, Metsada Pasmanik-Chor, Assaf Buch, Yuval Ebenstein, Paul Zimmet, Naftali Stern

## Abstract

**Background:** The epigenetic age can now be extrapolated from one of several epigenetic clocks, which are based on age-related changes in DNA methylation levels at specific multiple CpG sites. Accelerated aging, calculated from the discrepancy between the chronological age and the epigenetic age, has shown to predict morbidity and mortality rate. We assumed that deconvolution of epigenetic age to its components could shed light on the diversity of epigenetic, and by inference, on inter-individual variability in the causes of biological aging.

**Results:** Using the Horvath original epigenetic clock, we identified several CpG sites linked to distinct genes that quantitatively explain much of the interpersonal variability in epigenetic aging, with CpG sites related to secretagogin and malin being the most variable. We show that equal epigenetic age in different subjects can result from variable contribution size of the same CpG sites to the total epigenetic age. In a healthy cohort, the most variable CpG sites are responsible for accelerated and decelerated epigenetic aging, relative to chronological age.

**Conclusions:** Of the 353 CpG sites that form the basis for the Horvath epigenetic age, we have found the CpG sites that are responsible for accelerated and decelerated epigenetic aging in healthy subjects. However, the relative contribution of each site to aging varies between individuals, leading to variable personal aging patterns. Our findings pave the way to form personalized aging cards allowing the identification of specific genes related to CpG sites, as aging markers, and perhaps treatment of these targets in order to hinder undesirable age drifting.

## Background

In the past decade, the concept of epigenetic age has attracted growing interest and the number of publications on epigenetic clocks has risen exponentially (1), mostly since it appears to reflect at least some aspects of the biological age. The epigenetic age can be now extrapolated using one of several independently generated epigenetic clocks, each mathematically constructed from time/age related changes in DNA methylation levels at specific multiple CpG sites that collectively, with proper weighting, are highly correlated with chronological age (1–5). Discrepancies between chronological age and the calculated epigenetic are presumed to represent a measure of biological aging, such that epigenetic age acceleration /deceleration signifies accelerated or a relatively diminished rate of biological aging, respectively. Hence, epigenetic clocks can be compared to individuals’ chronological ages to assess inter-individual and/or inter-tissue variability in the rate of aging (6). How useful and informative this approach could be is exemplified by reports that epigenetic age is a predictor of time of death, mortality rate (2,4,7–9) and susceptibility to diseases such as lung cancer (10), breast cancer (11) and cardiovascular events (12).

Time/age related methylation appears to be a rather extensive process as is readily demonstrated by the fact that there are several different epigenetic clocks, each calculated based on several tens or hundreds different methylation sites which mostly do not overlap (13). Since these clocks vary with respect to their linkage to health outcomes, it is possible that each detects different processes which distinctly contribute to some facets of the biological age.

It is presently unknown whether the changes in methylation profiles which link epigenetic age acceleration to mortality and morbidity are simply aging markers or, perhaps, active players in the aging process. The implication of the latter is that reversal of epigenetic age could comprise a therapeutic target or at least, a measure of therapeutic success achieved by various pharmaceutical means (14) or perhaps lifestyle modification. For example, in murine studies, a reversal of the epigenetic age was achieved by reduced caloric intake (15). Assuming that the methylation level of the CpG sites, comprising the epigenetic clock, affect specific aging routs through modulation of gene expression, interpersonal differences in the methylation degree of such sites could offer clues not only to differential aging rates, but to variability in aging mechanisms in human subjects.

In the present study we focused on the possibility that the epigenetic age might be individually determined by inter-person differences in the methylation levels of such sites. For example, if the epigenetic age is kept fixed at Z years, which reflects the sum of three CpG sites, A, B and C, how variable is the specific contribution of each of them to the epigenetic age among different subjects? If A adds most years in one subject, but very little in another, might this reflect important differences in their aging driving mechanisms? What, if any, are the key epigenetic differences between “epigenetically young” and “epigenetically old” subjects of the same chronological age? This study, focuses on the interpersonal variability of the components (methylation levels at specific sites) comprising the epigenetic age, as a potential tool in predicting individual’s physiological malfunction, towards the development of personalized medicine. To address this issue we analyzed the epigenetic age of publicly available methylation data of 1441 healthy individuals, of the ages of 40-80 years, retrieved from Illumina methylation arrays. Because of the shorter lifespan of subjects with diabetes mellitus we have also analyzed datasets of 89 diabetic subjects. The epigenetic age was calculated by Horvath’s clock, which is based on coefficients, calculated by a regression model, relating the methylation status of 353 CpG sites (β-values) to chronological age.

## Results

### The epigenetic age distribution

In the aim of assessing interpersonal variability in the aging mechanisms, we first calculated the epigenetic age of the 1441 samples. Figure 1 depicts the overall relation between epigenetic age and chronological age in the entire analyzed data set. Individuals with an epigenetic age of Avg ± 1SD, (between red and orange lines), comprise 71.5% of the data whereas 27% of the data is from people with an epigenetic age between Avg ± 1SD and Avg ± 2.5 SD (between the orange and the purple lines). The 1.5% of the data residing beyond the purple lines were ignored to avoid large effects of potentially uncertain results (“outliers”). The near linear increase of the epigenetic age with chronological age, demonstrates that Horvath’s clock is suitable as age predictor for this data set, and that the epigenetic age has a high variability between individuals with the same chronological age, thus implying that individuals may age differently.

**Figure 1:**
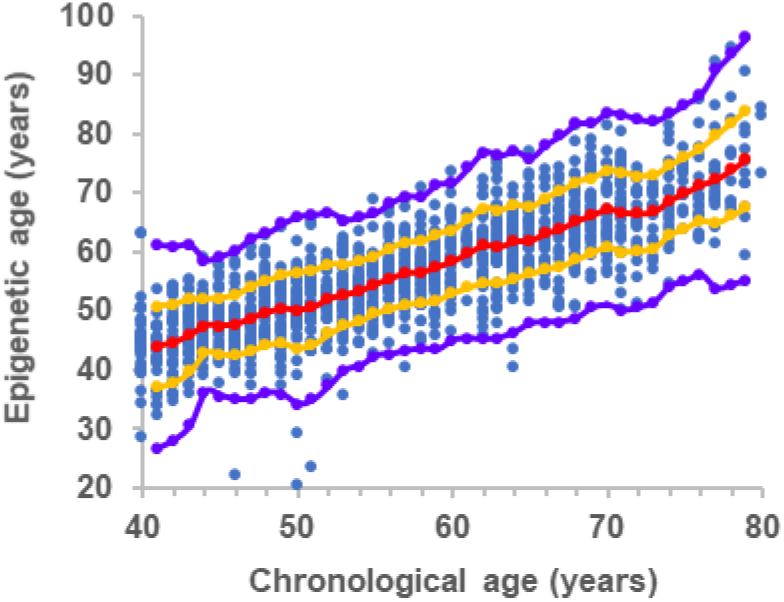
The overall relation between epigenetic age and chronological age. Each blue point represents a single healthy individual. Red dots are the average value of the epigenetic age at each chronological age connected by a regression curve (red line). All dots between the orange and the red line represent individuals with an epigenetic age between Avg and Avg ± 1SD. Dots residing between the orange and the purple lines represent individuals with an epigenetic age between Avg ± 1SD and Avg ± 2.5 SD.

### CpG sites with the highest inter-personal variability

In order to find out the possible cause for interpersonal variations in the epigenetic aging, we have searched for CpG sites that were the most variable among subjects, in terms of years added/subtracted by that site to/from the total epigenetic age. Epigenetic age-related DNA methylation is sex specific (8,16) as also shown in figure S1 in supplementary file 1. The β value of the components of the epigenetic age (Horvath’s 353 CpG sites) for the same chronological age group is also affected by sex, section 2.3 in supplementary file 1. Hence, we have examined whether the inter-personal variability of the components of the epigenetic age, is affected by sex, as well. Tables S2 and S3, in supplementary file 1 lists the CpG sites that reached the top 20 most inter-personal variable sites in 7 out of 8 chronological age groups, separately for men and women, according to the size of their average SD (in years, from top to bottom). We show that most of these top inter-personal variable CpG sites, were common for men and women, with very similar SD. We therefore decided to treat our data with no sex distinction and selected nine CpG sites that were most variable between individuals and were seen as prominently variable in both women and men (with similar SD) across the chronological age span of 40-80 years. These sites and the genes close to/ within which they are located are listed in table 1, ranked from the one showing the largest inter-personal variation (largest average SD of all chronological age groups) to the smallest. The two most dominant variable sites are related to secretagogin (SCGN) and malin [NHL Repeat Containing E3 Ubiquitin Protein Ligase 1 (NHLRC1)]. Eight probes out of the 9 most variable probes selected, were found to be independent of the population size (≥80% confidence). Specifically, secretagogin and malin were selected in 100% and 98% of the statistical simulations, respectively, indicating independence of the population size (table S4, in supplementary file 1).

**Table 1:** CpG sites with the highest inter-personal methylation variability in healthy population

| Related gene symbol | Related Gene definition/ product | Illumina's CpG ID | Contribution to epigenetic age (years)* | SD of the age contribution (years) |
| --- | --- | --- | --- | --- |
| SCGN | Secretagogin | cg06493994 | 7.1 | 2.1 |
| NHLRC1 | Malin | cg22736354 | 9.3 | 1.8 |
| MIR7-3HG | MIR7-3 Host Gene | cg02479575 | 1.8 | 1.0 |
| FZD9 | Frizzled 9 | cg20692569 | 6.8 | 0.9 |
| SCAP | SREBP cleavage-activating protein | cg26614073 | -4.7 | 0.7 |
| REEP1 | Receptor expression enhancing protein 1 | cg01968178 | 2.5 | 0.7 |
| CSNK1D | Casein kinase 1 delta | cg19761273 | -4.3 | 0.7 |
| FXN | Frataxin | cg07158339 | -4.6 | 0.7 |
| NDUFS5 | NADH dehydrogenase Fe-S protein 5 | cg07388493 | -4.8 | 0.7 |

*\*No sign for the contribution, in years, for each probe means a positive age contribution; a minus sign indicates a negative age contribution*

Notably, genes whose CpG sites showed the largest variation (SD) also contributed a sizable positive or negative age years to the calculated epigenetic age. However, inter-subject variability in methylation size effect was not simply a reflection of the magnitude of the contribution to age (in years). For example, the CpG site on type 1 hair keratin, protein phosphatase 1 regulatory (inhibitor) subunit 14/7 and the CpG site on testis expressed sequence 286 genes, which both have added/ subtracted more than +/-6.3 years to/ from the epigenetic age, did not add considerably, on average, to the interpersonal epigenetic age variability.

We have examined the correlation between the methylation levels of CpG sites with the highest inter-personal variability (from Horvath’s clock) and other CpG sites residing on the same gene. The level of the correlation was found to be related to the proximity of the CpG sites to one another and their location on the gene or its regulatory elements (figure S2 in supplementary file 1, and supplementary file 4). This indicates that the methylation level of the site included in the calculation of the epigenetic age provides good representation of the methylation status of neighboring sites and is therefore likely to be related to gene expression, if it resides on the promoter or another regulatory region.

### Interpersonal variations in the epigenetic age composition in subjects with identical epigenetic age

Quantitative variability in the aging vectors was not only found in individuals with identical chronological age and different epigenetic age but also in subjects with identical chronologic and epigenetic age. For example, figure 2 illustrates the heterogeneity of the contribution to the epigenetic age (in years) of the nine CpG sites that tend to vary the most among subjects. Figure 2A, shows two men, both at the chronological age group of 40-41, with a similar epigenetic age of 40-41. Despite these similarities, malin (NHLRC1) contributes more than 9.5 years to the epigenetic age in man P2 but only 7.5 years in man P1, a difference which is offset by larger contributions of at least two CpG sites-linked to age lowering, which are mapped on SCAP and FXN genes. Next, figure 2B shows two men at a chronological age of 40-41 years. Larger “aging” contribution of all major positive contributors, generated a larger cumulative aging effect in subject P3 compared to subject P4, amounting to a pro-aging effect of 9.5 years. Since both men have an epigenetic age of 45-46 years, this is offset, to some extent, by methylation state of CpG sites that lower the epigenetic age, particularly, in this case, sites mapped on the genes: CSNK1D FXN and SCAP. In the final example (figure 2C), two men of the same chronological age group of 40-41 have a markedly accelerated epigenetic aging of 50-51 years. However, age acceleration is driven by higher aging effect of CpG sites mapped to NHLRC1, SCGN and REEP1 in man P5, with a higher age-reducing effect of CSNK1D and SCAP, which only slightly make up for the stronger pro-aging CpG sites in this person. Presumably, the cumulative effect of other negative age contributors (that are less variable) are responsible for equalizing the epigenetic age in this later pair. This data implies that age acceleration at the same chronological age to the same higher or to the same equal epigenetic age can be reached by a highly variable methylation profile of the genes whose variation in general is the largest. This could signify inter-subject differences in the mechanisms underlying aging processes.

**Figure 2:**
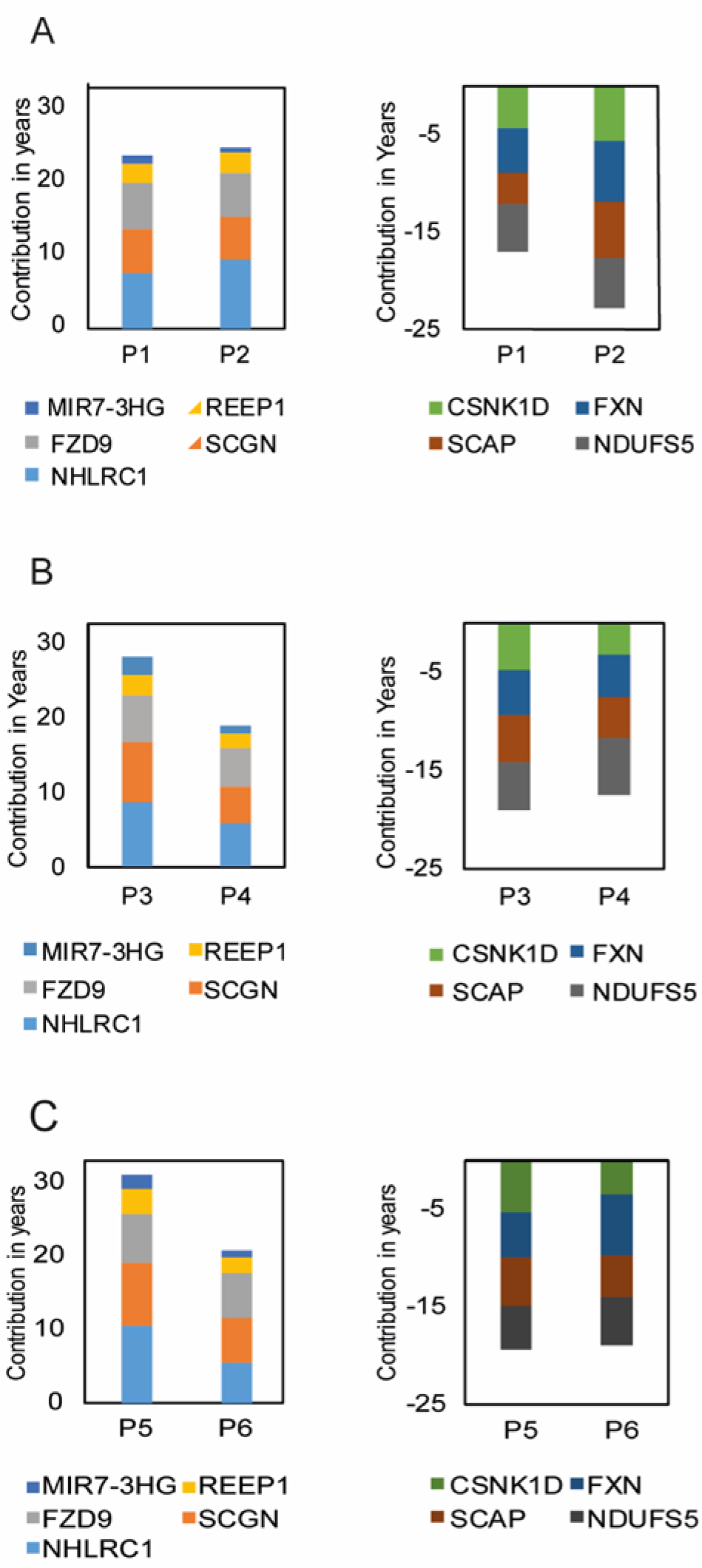
Interpersonal variability in the epigenetic age composition of individuals with the same chronological age of 40-41 years old and the same epigenetic age of (A) 40-41 years, (B) 45-46 years and (C) 50-51 years. Left and right panels are for CpG sites on genes which add or subtract from the epigenetic age, respectively. The genes presented according to the color codes at the bottom of each graph, are related to the CpG sites which belong to the nine sites with the highest variability

### Key CpG sites as age accelerators or decelerators

Figure 1 shows that almost a third (27%) of the individuals in our data set, spanning all chronological ages, and gender reside between Avg ± 1SD and Avg ± 2.5SD (dots between orange and purple lines). To detect which CpG sites (and their associated genes) were the major contributors to age acceleration we applied a “greedy algorithm” to the group of “epigenetic old” individuals (individuals whose epigenetic age resides between 1SD to 2.5SD, above the average epigenetic age line) and found that the site responsible for the largest “unfavorable aging” effect in 29% of the “epigenetically old” individuals, is mapped to the secretagogin gene (figure 3A). The remaining 70% of the “epigenetically old” individuals were then tested for the second largest ager, found to be malin. Once malin is consecutively normalized into the average zone, the epigenetic age of 12% of “epigenetically old” individuals is shifted to the average zone. This is followed by frataxin, responsible for 7%, and so on. Since frataxin is a negative age contributor, its effect is depicted by smaller age lowering vector. The entire group of genes related to the CpG sites, responsible for the accelerated aging of 80% of the healthy population, is presented in figure 3A. The CpG sites associated with secretagogin, malin MIR-7 and SCAP were found to be age accelerating components, independent of the population size (≥95% confidence, table S5 in supplementary file 1).

**Figure 3:**
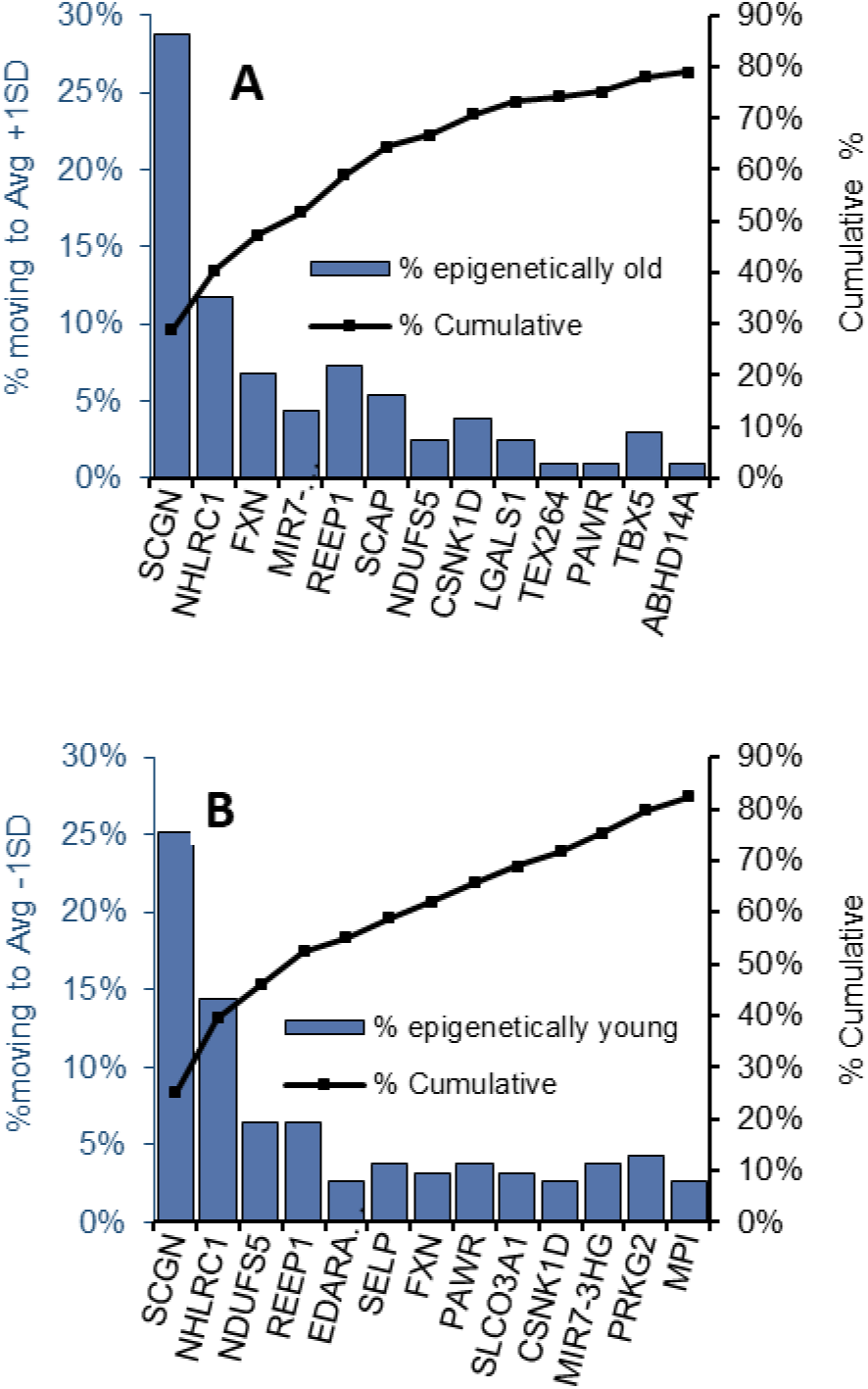
key genes associated with CpG sites, considered as age accelerators/ decelerators. (A) Blue bars present the percentage of individuals from the “epigenetically old” group which moved to the Avg + 1 SD group after setting consecutive CpG sites (presented with the name of their related gene) to their mean epigenetic age contribution, starting from the CpG site which moved the highest number of individuals to the lowest. (B) Blue bars are the percentage of individuals from the “epigenetically young” group who moved to the Avg - 1 SD after setting consecutive CpG sites to their mean epigenetic age contribution, starting from the site which moved the highest number of individuals to the lowest. The black line is the accumulative percentage of individuals moving from the “epigenetically old (A) /young (B)” group to the average group.

The same key players, with some change in the magnitude and order of their effect and some new effectors’, stood out when we searched for the genes associated with the CpG sites responsible for lowering the epigenetic age (age decelerators) from the Avg - 1SD age zone to an epigenetic age of less than -1SD down to – 2.5 SD (figure. 3B). Secretagogin and malin were the largest contributors to age deceleration, accounting for 25% and 14% of the “epigenetically young” individuals, respectively. The cumulative percentage of individuals moving from the epigenetic old/young group to the Avg ± 1SD (black line in figure 3A and B) shows that up to 80% of the epigenetically old/young individuals had a single key prominent gene, responsible for accelerated or decelerated aging.

Collectively, then, many of the CpG sites or their associated genes that are responsible for interpersonal variation in the makeup of the epigenetic age, are also major players that act as “age accelerators” or “decelerators”, depending on their methylation status. The fact that the same two CpG sites, residing near the genes secretagogin and malin, are responsible for the accelerated aging of 41% of the population and for the decelerating aging of 39% of the population (i.e the same two CpG sites are responsible for the fact that 27% of individuals in our data set, reside between Avg ± 1SD and Avg ± 2.5SD), implies that the key CpG sites responsible for individuals residing between average ± 1SD and average ± 2SD of the epigenetic age are not random or due to noise fluctuation. In addition, we show in our bootstrapping simulations, that these two genes are age accelerators/ decelerators with above 98% confidence! (appearing in above 98% of the simulations), implying no coincidence in the selection of these key components (table S5, supplementary file 1).

### Personalized epigenetic aging card

Finally, a putative “personal epigenetic aging status card” can be produced for each individual tested by the Horvath epigenetic clock. The aging card is based on the methylation status of the nine CpG sites, that were found to consistently contribute more than others to the interpersonal variability in both the men and women cohorts and over eight age groups spanning 40 to 80 years. As such, this card is adequate for the aging analysis of both sexes and to individuals at the age range of 40 to 80 years. As shown in figure 4 for 7 individuals, all at the same chronological age of 40-41, this card grades each subject for the accelerating (orange to red cells, figure 4) or decelerating effect (green cells, figure 4), in years of each of the nine most variable CpG sites. The grades are the relative deviation of the age contribution of each CpG site from its average contribution to the epigenetic age. If the difference between the calculated epigenetic age of a certain individual and the average epigenetic age cannot be significantly explained by these sites, the clock can be further interrogated to reveal other sites, with less common age effect, which might explain deviations from the average epigenetic age. This process may eventually evolve as an individualized panel of aging effects, much like a routine biochemistry panel as presently assessed at the clinician’s office to detect indicators of disease, by their actual deviation from the normal range.

**Figure 4:**
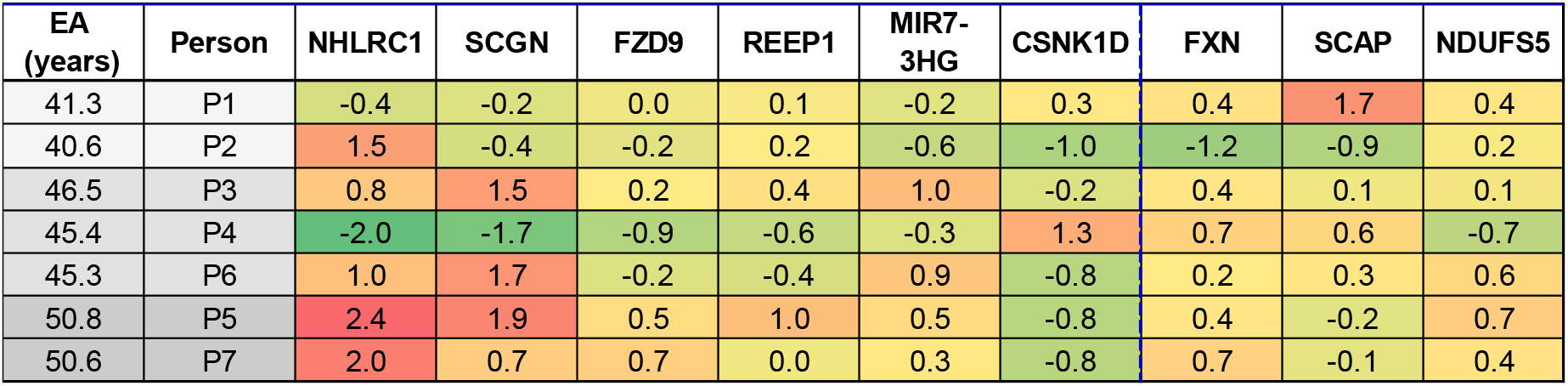
Personalized epigenetic gene card: colored cells are the deviation of the epigenetic age contribution (in years) of the CpG sites or their associated genes from their average epigenetic age contribution, for seven individuals at chronological age of 40-41 years (sample number is # GSM). The average epigenetic age for 40-41 years old men, is 45.7 years. We show two samples ~5 years below epigenetic average age (light gray), three samples at average epigenetic age (darker gray) and two samples ~5 years above epigenetic average age (dark gray). Cells with orange to red colors are for genes associated with CpG sites with age contribution above average. Cells in green, or light green associate with CpG sites on genes with age contribution below average.

### Epigenetic aging variability in diabetic mellitus

In order to find the CpG sites responsible for the variability in the epigenetic aging of diabetic subjects, we first calculated their epigenetic age, using Horvath’s epigenetic clock. Our diabetic dataset consisted of 89 subjects of which 63 had type 1 diabetes (T1D) and 26 had type 2 diabetes (T2D) at the age ranges of 40-70 years and 65-80 years, respectivaly. Figure 5 depicts the calculated epigenetic age of the diabetic subjects as a function of their chronological age, superimposed on the epigenetic age graph, presented for the healthy population in figure 1. According to figure 5, the epigenetic age of the T1D subjects (red and black triangle marks, figure 5) was lower than the average epigenetic age of the healthy population (bellow the red diagonal circles in figure 5) and the epigenetic age of the T2D subjects (green triangle marks, figure 5) was around the average epigenetic age of the healthy population (around the red diagonal circles in figure 5). In addition, no differences were observed in the average epigenetic age of T1D subjects receiving intensive treatment and showing no complications (red triangle) and those on conventional therapy who also developed albuminuria and/ or diabetic retinopathy (black triangle).

**Figure 5:**
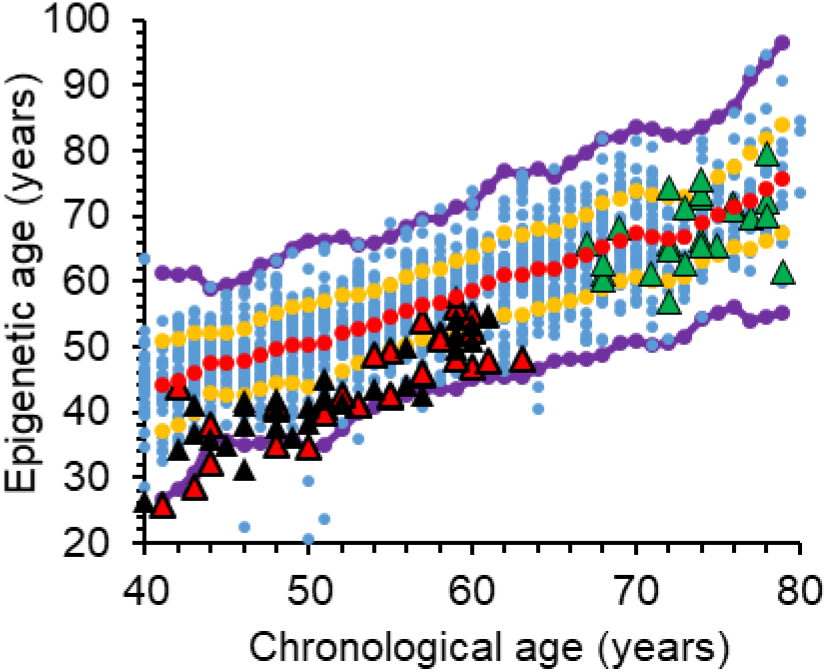
The overall relation between epigenetic age and chronological age, including the samples from diabetic subjects. Each blue point represents a single healthy individual. Red dots are the average value of the epigenetic age of the healthy cohort at each chronological age. The diabetic subjects are represented by triangles, superimposed on the general healthy cohort; red triangles are for T1D subjects receiving intensive treatment and are free of complications; black triangle are for T1D subjects receiving conventional therapy who developed microvascular complications (albuminuria and /or retinopathy); and green triangle marks are for T2D. All dots between the orange and the red line represent individuals with an epigenetic age between the average and average ± 1SD. Dots residing between the orange and the purple lines represent individuals with an epigenetic age between the average ± 1SD and the average ± 2.5 SD.

The CpG sites which contributed the largest number of years to the interpersonal variations in the epigenetic age of the diabetic subjects (highest SD, in years) were first derived separately for the T1D and T2D data sets. The 20 CpG sites that showed the largest inter- personal variability that were consistently present in all 5 and 3 chronological age groups, in the T1D and T2D datasets, respectively are listed in tables S6 and S7, in supplementary file 1. Since the leading 5 most variable CpG sites were common for both T1D and T2D, we combined these datasets and treated them as a diabetic data set so as to cover the entire studied range of chronological ages with a better statistical power. The results of the analysis of the combined file, including the ranked list of the most variable sites, the magnitude of their variability (i.e., SDs) and their associated genes are provided in table 2. As in the healthy population, the two most variable sites in the diabetic dataset were related to malin (NHLRC1) and secretagogin (SCGN). Frizzled 9 (FZD9) also reached the top 5 sites both in the healthy and in the diabetic cohorts. However, two new CpG sites that are related to PRKC Apoptosis WT1 Regulator (PAWR) and L-pipecolic acid oxidase (PIPOX) genes emerged at the top most variable sites only in the diabetic cohort. Interestingly, in the healthy population the site related to PAWR gene appears among the 20 CpG most variable sites only in the age groups of 55 years or higher (data not shown), whereas, in the diabetic population the PAWR site appears as the third most variable gene in all age groups.

**Table 2:** CpG sites with the highest inter-personal methylation variability in the combined T1D + T2D cohort

| Related gene symbol | Related Gene definition/ product | Illumina's CpG ID | Contribution to epigenetic age (years)* | SD of the age contribution (years) |
| --- | --- | --- | --- | --- |
| NHLRC1 | Malin | cg22736354 | 9.6 | 1.7 |
| SCGN | Secretagogin | cg06493994 | 7.6 | 1.4 |
| PAWR | PRKC Apoptosis WT1 Regulator | cg00864867 | 3.2 | 1.2 |
| PIPOX | L-pipecolic acid oxidase | cg06144905 | 3.3 | 0.7 |
| FZD9 | Frizzled 9 | cg20692569 | 6.6 | 0.6 |

## Discussion

Following the pioneering reports by Horvath (1) and Hannum (5), it is now widely recognized that epigenetic age and chronological age correlate well with each other in many populations, regardless of the tissue studied (1). In several publications a potential functional role has been ascribed to the CpG sites that participate in the algorithms developed to derive the epigenetic age (3). For example, accelerated epigenetic age relative to chronological age is reportedly linked to preferential activation of pro-inflammatory and interferon pathways, along with reduced stimulation of transcriptional/translational machinery, blunted DNA damage response, and weakened mitochondrial signatures (3),(17). This supports the notion that the epigenetic age does not simply mirror randomly the passage of time but reflects specific anti-homeostatic effects that may lead to or indicate specific unfavorable conditions which facilitate disease and affect life span.

The possibility that specific epigenetic aging drivers can be targeted to achieve personalized epigenetic/biological age deceleration, can become a testable approach. In the present study we examined which of the CpG sites included in the original Horvath DNAm algorithm are the major contributors to inter-personal differences in the epigenetic age. We found that the CpG sites related to malin and secretagogin have relatively high contribution to the epigenetic age and the most variable methylation status in between individuals both in the healthy and the diabetes cohorts. Additionally, in the healthy cohort, differences in the methylation status of secretagogin and malin, contributed more than any other methylation loci, to the difference between epigenetically old and epigenetically young subjects, across the entire age span screened by us (40-80yrs) (figure 3).

Notably, both the secretagogin and malin-related CpG sites are among the 5 shared by three known DNA methylation -based epigenetic clocks: Hannum DNAm Age score (based on 71 methylation sites) (5), Horvath DNAm Age measure (353 sites) (1) and the DNAm PhenoAge score (513 sites) (2) and were also suggested to be the most dominant key age predictor sites (18). Might these genes, then, be mechanistically involved in aging?

The malin gene encodes a RING type E3-ubiquitin ligase which forms a functional complex with laforin, a glucan phosphatase (19). Mutations in either malin or laforin in humans lead to the development of Lafora progressive myoclonus epilepsy, a rare fatal neurodegenerative disease with early manifestations in the early childhood. Brain damage is incurred due to deposition underbranched and hyperphosphorylated insoluble glycogen in the brain and peripheral tissues (20–22). It is notable that glucan deposits have been described in the setting of aging animals and humans (23–25), unrelated to LaFora disease, which raises the possibility of lesser malin activity with age. Indeed, malin appears to participate in a delicate homeostatic network linking neuronal glycogen synthesis and energetic utilization, interacting with autophagy, mitochondrial function and response to thermal stress, which could collectively affect lifespan (19), (25–28). The possibility that malin expression, which is critical for inhibition of polyglucan deposits in neurons, plays a role in healthful longevity in humans is intriguing and requires targeted research. In animal studies malin deficiency can lead to impaired autophagy and accumulation of dysfunctional mitochondria, which eventually promote neurodegeneration, immune disorders, cancer, and accelerated aging (27).

Secretagogin is an intracellular calcium sensor and facilitator of insulin secretion by pancreatic islet beta cells (29). Recently it was shown that secretagogin play a critical role in the second phase of glucose-stimulated insulin secretion (30), protect against insulin aggregation and enhance peripheral response to insulin (31). Concordant with this broad role in carbohydrate handling, secretagogin knockout leads to hyperglycemia (32). Secretagogin is also expressed in neuroendocrine cells where it likely regulates exocytosis and hormone release (33,34). Concordantly, it is also involved in danger avoidance behavior through the control of post synaptic cell-surface availability of NMDA receptors in the central amygdala (35). We are not aware, however, of published reports examining the relation between induced changes in secretagogin expression and lifespan or longevity.

Of major interest in the Horvath algorithm are CpG sites with a negative contribution to the epigenetic age, such as frataxin. Frataxin is a nuclear-encoded mitochondrial protein which is part of the Fe-S-cluster-containing proteins acting as an iron chaperone, thereby allowing normal function of the mitochondrial respiratory chain (36). In our analysis frataxin shows both high interpersonal variability and also partly explains some (~8%) of the calculated age difference between epigenetically old and average subjects (figure 3A). The fact that higher methylation of frataxin can extend life, as indirectly suggested by its epigenetic age lowering effect is somewhat counterintuitive: defects in the expression of this mitochondrial protein cause the neurodegenerative syndrome of Friedreich’s ataxia (37,38), which is also accompanied by cardiomyopathy, diabetes mellitus and reduced life expectancy (39). However, inactivation of many mitochondrial genes in the nematode Caenorhabditis elegans by RNAi was actually shown to extend lifespan (40). Ventura et al reported that suppression of the frataxin homolog gene (frh-1) prolonged lifespan in the nematode, along with an altered phenotype of smaller size, diminished fertility and variant responses to oxidative stress. Thus, whereas sizable inactivation of frataxin causes a disabling disease, a more moderate frataxin suppression, such as achieved by RNAi could lead to higher lifespan as seen in C. elegans (41). There is evidence that frataxin silencing induces mitochondrial autophagy as an evolutionarily conserved response to the ensuing iron starvation (36). In a broader sense, lesser frataxin availability might comprise a surmountable challenge which elicits mitophagy that eventually preconditions the cell’s capacity to sustain future stress, thereby increasing the likelihood of extended lifespan.

Our findings of inter-personal variabilities in the epigenetic age components of the healthy population, has raised our interest in discovering epigenetic age patterns of individuals with biological-age accellerating diseases, such as, diabetic patients. Our results did not reflect epigenetic age acceleration for T2D and showed rather the oposite, for T1D subjects which had lower epigenetic age than the average of the healthy population (under the red curve in figure 5). Nevertheless, these results are in agreement with earlier publications which have also used chronological age-based epigenetic age calculators, such as Horvath’s epigenetic clock (8,42). This may indicate that the CpG probes chosen for the construction of such epigenetic age clocks, do not reflect variations in DNA methylation leading to epigenetic age drifting in diabetes or its complications. What makes the T1D population “epigeneticly younger” according to the aging clock used in this study, would be an interesting question for future investigations. Notably, a different epigenetic clock type, the “DNAm GrimAge”, which incorporates DNA methylation sites related to surrogate biomarkers of smoking level and of selected plasma proteins, that are strongly associated with mortality and morbidity may be a better choice for predicting age acceleration in diabetics (3,4,42).

Of the top five most variable age components that are found solely in the diabetic cohort, PAWR is a tumor suppressor gene, inducing selective apoptosis of cancer cells (43–45), and may thus be related to association of aging with higher cancer rates. Another component is related to PIPOX which catalyzes the oxidation of L-pipecolate, an intermediate step in the catabolic process of L-lysin to acetyl-CoA, produced in the pipecolate pathway (46,47). Elevated levels of lysine were found to be associated with higher risk for the development of T2D and for T2D-concomitant cardiovascular disease (CVD) (48). In addition, PIPOX promotes sarcosine oxidative N-demethylation, yielding glycine (49), as part of the sarcosine pathway, which is involved in the methionine cycle (50–53). The methionine cycle is responsible for the production of S-adenosylmethionine (SAM), the methyl donor substrate in the process of cytosine DNA methylation by the family of DNA methyl transferase (DNMT) enzymes. Differential levels of PIPOX and sarcosine were observed in several types of cancers (49,54). In addition, methionine cycle restriction and regulation of SAM production was shown to extend lifespan in various animal models (55).

## Conclusions

Overall, our analysis reveals sizable interpersonal differences in the contribution to age of methylation sites of several genes. It is also possible that there is a shift in the epigenetic age vectors in diabetes mellitus patients that are not necessarily detected by the computation of the mean epigenetic age per se. In the healthy cohort, genes such as, but not limited to, secretagogin, malin and frataxin stand out in terms of either the size of their effect on interpersonal differences in the composition of the epigenetic age as well as their influence on the likelihood for an individual to acquire enhanced or delayed epigenetic aging. This analysis also unravels that even healthy subjects with average epigenetic aging could show accelerated aging with respect to some genes. In the same venue, epigenetically healthy older subjects are also heterogeneous and could be pushed to unfavorable epigenetic drifting by different aging vectors. Interestingly, malin and secretagogin were also found to be the most variable age components in diabetic subjects. Two additional age components, PIPOX and PAWR, entered the top 5 most variable sites, only in diabetic patients. This paves the way for future attempts to personalize the perception of epigenetic aging by deconvolution, addressing aging not as a general process in search of reversal, but as a collection of individual effects requiring personalized attention.

## Materials and Methods

### Methylation Data

The β values which reflects the methylation status of each CpG site were retrieved from the Gene Expression Omnibus (GEO) Datasets repository. Following filtration for whole blood and healthy subjects at the age range of 40-80 years old, we obtained 2298 samples, from 23 different data sets. All filtered samples were normalized using an R-code provided by Horvath et al. [2]. Samples that failed Horvath’s normalization process (accounting for the two different designs of the methylation arrays, Type-I and Type II) [2], were removed, leaving a total of 1,441 samples for interrogation, (867 females and 574 males). We also analyzed two other distinct cohorts of diabetic subjects (n=89) with whole blood samples from the same age range, diagnosed with diabetes (63 T1D, age range 40-65 years old, 31 women and 32 men and 26 T2D, age range 65-80 years, 11 women and 15 men), a disease possibly linked to accelerated biological aging (56–58). Detailed description on the datasets and the processes involved in the selection of the samples we have analyzed can be found in table S1, detailed materials and methods section, in supplementary file 1.

### Epigenetic vs Chronological age

For each sample we extracted the β values of Horvath’s 353 CpGs clock sites and converted them to age contribution based on their coefficients, calculated as explained by Horvath et al. [2]. The epigenetic age of each of the 1,441 individuals is the sum of the contribution plus a constant (representing the intercept of the linear correlation), in years, of all 353 CpGs (supplementary file 2). In order to smooth the average (Avg) and standard deviation (SD), we used a running average and a running standard deviation of the epigenetic age with a window size of 3 (details in materials and methods section of supplementary file 1).

### CpG sites with the highest inter-personal variability

The 1441 healthy samples were first divided according to gender. Each gender group was further divided to 8 chronological data sets by age groups spanning from the age of 40 to the age of 80 years. At each chronological data set, we recorded 20 CpG sites from a total of 353 CpG sites from Horvath’s clock, with the highest inter-individual variability calculated as average standard deviation (average SD, in years). The average SD was calculated separately for each chronological age group and was expressed in years. Out of these outstanding 20 CpG sites we then identified the CpG sites which were also consistently present at least in 7 chronological age data sets, for each sex (table S2 and S3 in supplementary file 1). Most of the top CpG sites responsible for the variability within the population, across all chronological age data sets, were common to both men and women. We therefore decided to treat the data with no sex distinction. As a result, the entire data set of 1441 healthy samples was re-divided to 8 chronological data sets by age groups spanning from the age of 40 to the age of 80 years regardless of the sex (40-44 (189); 45-49 (215); 50-54 (217); 55-59 (223); 60-664 (220); 65-69 (177); 70-74 (120); 75-80 (80), years (number of subjects)).

A similar process was applied for the diabetic cohort; we first looked for the most variable CpG sites in the datasets of T1D and T2D, separately. We divided each dataset to chronological age groups with five years’ intervals: 5 chronological age groups spanning 40 to 65 years and 3 chronological age groups spanning 65 to 80 years, for T1D and T2D, respectively. We have then recorded the 20 CpG sites with the highest inter-individual variability (average SD, in years) for each age group within each dataset. We have found, for each dataset, 8 CpG sites which were consistently present in at list 5 and 3 chronological age groups in the T1D and T2D, respectively (table S6 and S7 in supplementary file 1). The 5 most variable CpG sites out of the 8 selected, across all chronological age groups, for each data set, were common to both T1D and T2D with slight differences in the magnitude of their variability (average SD, in years). Based on these results, we combined the T1D and T2D datasets as a consequence all age groups were covered in a single unified dataset so as to allow a more valid analysis (due to larger number of subjects in the dataset). We then divided the combined diabetic data set of total 89 subjects to 8 chronological age groups with 5 years intervals, spanning from 40 to 80 years, (40-44 (10); 45-49 (12); 50-54 (14); 55-59 (17); 60-664 (9); 65-69 (5); 70-74 (12); 75-80 (8), years (number of subjects)).

Finally, in order to find the most variable CpG sites in both healthy and diabetic data sets, we recorded the 20 CpG sites from Horvath’s clock, with the highest inter-individual variability (standard deviation, SD, in years), at each chronological age group. We then identified 9 and 5 CpG sites out of these 20 CpG sites that were consistently present in each of the 8 chronological age groups of the healthy and diabetic cohorts, respectively. A statistical simulation, examining the relation between the identified CpG sites and the size of the cohort was manifested as explained in supplementary file 1.

### CpG sites as age accelerators or decelerators

Key epigenetic age accelerators or decelerators were found by looking for the probes with the highest cumulative contribution to the epigenetic age. The data set was divided to: 1) the “epigenetically average” group including all samples with epigenetically age of the running average ± 1 SD (the population in between the two orange lines), 2) the “epigenetically old/young” group, with an epigenetic age between 1 SD and 2.5 SD above or below the average (the population in between the upper/lower orange and purple lines, respectively), 3) the outliers, which have an epigenetic age with more than 2.5 SD from the average (the population above the upper or below the lower purple lines).

For the “epigenetically old” and the “epigenetically young” population, a greedy algorithm was applied. The algorithm calculates, for each probe, the number of individuals that moved from the “epigenetically old” or the “epigenetically young” to the “epigenetically average” group, as a result of setting a particular probe to its mean epigenetic age contribution value (in years). In each iteration, the algorithm selects the probe which moves the largest number of samples into the “average group”: In the first iteration, the CpG site selected is the one which moves the highest number of subjects into the average zone, (by setting it to its average value). In the second iteration, the probe selected is the one which moves the most individuals in the residual “epigenetic older/ younger” zone to the average epigenetic age group and so on. The bars in the graph shown in figure 3 presents the percentages of individuals from the entire 1441 population, passing from the “epigenetically old/ young” to the average group when all consecutive CpG sites are set to their mean epigenetic age contribution (supplementary file 1)

### Personalized epigenetic aging card

A personal epigenetic card is presented for 7 healthy individuals, with chronological age of 40-41 years, as the deviation (in years) from the mean epigenetic age contribution of each of the 9 chosen probes. The mean epigenetic age contribution of each probe is the average addition/subtraction of each probe, to/ from the average epigenetic age at 40-41 years.

## Supplementary Materials

Supplementary File 1: 1. Detailed materials and method: 1.1. Description of methylation datasets and their filtration processes, Table S1: methylation Datasets of healthy and diabetic subjects. 1.2. An R code for the conversion of idat values β values. 1.3. Epigenetic age vs. chronological age. 1.4. CpG site cluster maps. 1.5. CpG sites as age decelerators or accelerators. 1.6. Statistical analysis. 2. Supplementary figures and tables: 2.1. Figure S1, The average epigenetic age of men is approximately 2 years higher than that of women. 2.2. Table S2: Gender-related differences in the methylation status of Horvath’s 353 CpG sites within the same age group, by one way ANOVA analysis. 2.3. Table S3: Gender specific CpG sites with the highest inter-personal methylation variability. 2.4. Figure S2: Correlation of methylation level between each selected CpG site and proximate CpG sites. 2.5. Table S4: The dependency of population size on the selection of the most interpersonal variable CpG sites, by statistical simulation. 2.6. Table S5: Statistical simulation for the dependency of the CpG sites chosen by the greedy algorism and population size. 2.7. Table S6: CpG sites with the highest inter-personal methylation variability in T2D cohort, Table S7: CpG sites with the highest inter-personal methylation variability in T1D cohort, Table S8: A comparison between CpG sites with the highest inter-personal methylation variability in T1D, T1D, T1D+T2D and healthy cohorts.

Supplementary file 2: The age contribution, in years, for each of the 353 CpG sites comprising Horvath’s clock, for each of the 1441 individuals

Supplementary file 3: The SD and the rank by SD of the most variable CpG sites from Horvath’s clock in three different populations: Men, Woman and gender mixed population

Supplementary file 4: The correlation between the methylation of the CpG site included in Horvath clock and other CpG sites related to the same gene, as a function of their distance, for the nine selected most variable genes found in all age groups.

## Ethics approval and consent to participate

The study has received ethics approval from the Tel-Aviv Sourasky Medical Center Institutional Review Board. Informed consent was obtained from all subjects involved in the study.

## Consent for publication

Not applicable.

## Availability of data and materials

The GEO number of the datasets analyzed during the current study are listed in supplementary file 1, section 1, table S1. The data generated during the current study is available from the corresponding author on reasonable request

## Author Contributions

N.S. initiated the project. N.S., E.S., T.K and T.S. designed the study. T.K, E.S, T.S, Y.M, G.S, and A.B have selected and filtered the data source. T.K and E.S did all the mathematical and statistical calculations. Y.E, P.Z and M.P gave useful reviews and comments on the paper. N.S and T.S. wrote the manuscript. All authors gave approval to the final version of the manuscript.

## Acknowledgments

We would like to thank Sami and Tova Sagol for their generous donation to the Sagol Epigenetic center, which made this study possible.

