## Supplementary file 1 for "Deconvolution of the epigenetic age discloses distinct inter-personal variability in epigenetic aging patterns"

**for the article**

### Contents

1. Detailed materials and methods

1.1. Description of methylation datasets and their filtration processes……..…..2

Table S1: methylation Datasets of healthy and diabetic subjects…………3

1.2. An R code for the conversion of idat values β values…………………………3

1.3. Epigenetic age vs. chronological age………………………………………..…4

1.4. CpG site cluster maps……………………………………………………….....4

1.5. CpG sites as age decelerators or accelerators……………………………..…5

1.6. Statistical analysis…………………………………………………………...…5

2. Supplementary figures and tables…………………………..………………......6

2.1. Figure S1: Gender effects on epigenetic age............................………………6

2.2. Table S2 and S3: Gender specific CpG sites with the highest inter-personal methylation variability…………………………………………………………..6

2.3. Gender differences in the β values of the nine selected CpG sites ………..…..8

2.4. Figure S2: Correlation of methylation level between each selected CpG site and proximate CpG sites……………….……………….………………...…...8

2.5. Table S4: The dependency of population size on the selection of the most interpersonal variable CpG sites, by statistical simulation……………………10

2.6. Table S5: Statistical simulation for the dependency of the CpG sites chosen by the greedy algorism and population size……………………………...……10

2.7. Tables S6, S7 and S8: CpG sites with the highest inter-personal methylation variability in the diabetic and the healthy cohorts………………….........…12

### 1. Detailed Materials and methods

#### 1.1. Description of methylation datasets and their filtration processes

Data was retrieved from the Gene Expression Omnibus (GEO) Datasets repository (https://doi.org/10.1093/nar/gks1193), up to august 2020, from three Illumina BeadChip platforms: GPL8490, GPL13534 and GPL21145 (Illumina HumanMethylation27 BeadChip, Illumina HumanMethylation450 BeadChip, Infinium Methylation EPIC, respectively). The β value which reflects the methylation status of each CpG site, derived from the relative fluorescent intensities of the methylated sites and the total (methylated + unmethylated) sites, was downloaded for a total of 134,942 samples. Samples with idat values were converted to β values using an R-code as described below. In the aim of working only with whole blood samples from heathy population we carried out a two- stage filtration: Initially, we automatically filtered out 123,529 samples with either missing data regarding the chronological age, gender, or that did not include the term "blood" in the description. Samples which were from individuals younger than 40 or older than 80 were excluded, as well. The description and the reference of the remaining 11,413 samples were manually selected for whole blood and healthy state, resulting in 2298 samples, from 23 different data sets (supplementary file 2). All filtered samples were then normalized for platform/bead type/ batch/ array using an R-code provided by Horvath et al. [1]. Samples that failed during the Horvath's normalization process [1] were removed, thus leaving a total of 1,441 samples for interrogation, of which 867 were females and are 574 males. We also analyzed a distinct cohort of 89 whole blood samples from subjects of the same age range, diagnosed with diabetes, a disease possibly linked to accelerated biological aging [2]–[5]. A list of all the datasets used in our analysis is presented in table S1.

Table S1: methylation Datasets of healthy and diabetic subjects

| Healthy | | Diabetes | |
|---|---|---|---|
| GSE | Platform | GSE | Platform |
| GSE87571 | GPL13534 | GSE76169 | GPL16304 |
| GSE111629 | GPL13534 | GSE62003 | GPL13534 |
| GSE50660 | GPL13534 | | |
| GSE41037 | GPL8490 | | |
| GSE58045 | GPL8490 | | |
| GSE106648 | GPL13534 | | |
| GSE20236 | GPL8490 | | |
| GSE64495 | GPL13534 | | |
| GSE67751 | GPL13534 | | |
| GSE52588 | GPL13534 | | |
| GSE32396 | GPL8490 | | |
| GSE85506 | GPL13534 | | |
| GSE77445 | GPL13534 | | |
| GSE53128 | GPL13534 | | |
| GSE40005 | GPL13534 | | |
| GSE52113 | GPL13534 | | |
| GSE107143 | GPL13534 | | |
| GSE41169 | GPL13534 | | |
| GSE99624 | GPL13534 | | |
| GSE32148 | GPL13534 | | |
| GSE120307 | GPL13534 | | |
| GSE100825 | GPL21145 | | |
| GSE84003 | GPL13534 | | |
| GSE111165 | GPL13534 | | |

1.2. An R code for the conversion of idat values β values:

```
#//package installation process

#if (!requireNamespace("BiocManager", quietly = TRUE))
 # install.packages("BiocManager")

BiocManager::install("ChAMP")
BiocManager::install("DMRcate")

library("ChAMP")

setwd("/Users/thomas/Desktop/Hidat/850K")
```

```
import <- champ.import(directory = getwd(), offset=100,
arraytype="450k")

write.csv(import$beta,"/Users/thomas/Desktop/Hidat/850K/850K_i
dat_parsed.csv")
```

1.3. Epigenetic vs Chronological age:

The epigenetic age was calculated by Horvath's clock, which relates the methylation status of 353 CpG sites (β- values) to chronological age [1]. For each sample we extracted the β values of these 353 CpGs sites and converted them to age contribution based on their coefficients calculated as explained by Horvath et al. ([1], supplementary file 3). A positive coefficient means that the methylation of the CpG site increases with age and a negative coefficient means that the methylation of the CpG site decreases with age. The epigenetic age of each of the 1,441 individuals is the sum of the contribution plus a constant (representing the intercept of the linear correlation), in years, of all 353 CpGs.

The epigenetic age was plotted against the chronological age given in the sample description. In order to smooth the average (Avg) and standard deviation (SD), we used a running average and a running standard deviation of the epigenetic age with a window size of 3, meaning that each point in the running average graph is the average for all samples at a particular age and the samples of one year above and one year below that age. In the same manner the running SD is the SD of the samples at a certain age ±1 year. The red dots in figure S1 (main paper) are the running average value of the epigenetic age connected by a red line. The "ORANGE" lines represent the running average ±1 running SD (Avg ± 1SD) and the "PURPLE" lines represent the average ± 2.5 running SD (Avg ± 2.5SD).

1.4. CpG site cluster map

For the creation of cluster maps, each of the 9 CpG sites that were the most variable in all 8 healthy data sets, was compared with other Illumina's 450K CpG sites related to the same gene. The list of CpG sites related to each gene was taken from Illumina's website. The Pearson correlation coefficient between each pair of probes based on the 1441 individual CpG sites was calculated creating the correlation matrix. These probes were clustered based on Euclidean distance using hierarchical clustering and the correlation matrix rows and columns were arranged based on the resulting clusters.

1.5. <u>CpG sites as age accelerators or decelerators</u>

Key epigenetic age accelerators or decelerators were found by looking for the probes with the highest cumulative contribution to the epigenetic age. The entire data set was divided to four groups of samples: 1) the "epigenetically average" group including all samples with epigenetically age of the running average ± 1 SD (the population in between the two orange lines), 2) the "epigenetically old" population, which are those who have an epigenetic age between 1 SD and 2.5 SD above the average (the population in between the upper orange and purple lines), 3) the "epigenetically young" population, which are those who have an epigenetic age lower than 1 SD but not less than 2.5 SD below the average (the population in between the lower orange and purple lines), 4) the outliers, which have an epigenetic age with more than 2.5 SD from the average (the population above the upper purple or below the lower purple line).

For the "epigenetically old" and the "epigenetically young" population, a greedy algorithm was applied. The algorithm calculates, for each probe, the number of individuals that moved from the "epigenetically old" or the "epigenetically young" to the "epigenetically average" group, as a result of setting a particular probe to its mean epigenetic age contribution value (in years). In each iteration, the algorithm selects the probe which moves the largest number of samples into the "average group": In the first iteration, the CpG site selected is the one which moves the highest number of subjects into the average zone, (by setting it to its average value). In the second iteration, the probe selected is the one which moves the largest number of individuals in the residual "epigenetic older/ younger" zone to the average epigenetic age group and so on. The bars in the graphs shown in figure 4 presents the percentages of individuals from the entire 1441 population, passing from the "epigenetically old/ young" to the average group when all **consecutive** CpG sites are set to their mean epigenetic age contribution.

1.6 <u>Statistical analysis</u>

In order to check whether our results may be influenced by the population size, we have conducted a statistical analysis which was based on 1000 bootstrapping simulations [6], [7]. The bootstrapping <u>is a statistical procedure that resamples a single dataset to create many simulated samples</u> allowing the calculation of confidence intervals, p-val and any statistical measurements. The population size, parameter and the cohort chosen for the simulation are dependent on the relevant study.

### 2. Supplementary figures and tables

#### 2.1. Figure S1: Gender effects on epigenetic age

Age related DNA methylation is gender specific [8], [9]. Our initial analysis was conducted on men and women separately. For example, we have found that, in a population of 40-41 years old individuals, the epigenetic age of men, on average, is 5.4 years older than their chronological age whereas women, on average, are only 2.6 years older than their chronological age. Hence, men, on average, have an additional 2 years of accelerated aging.

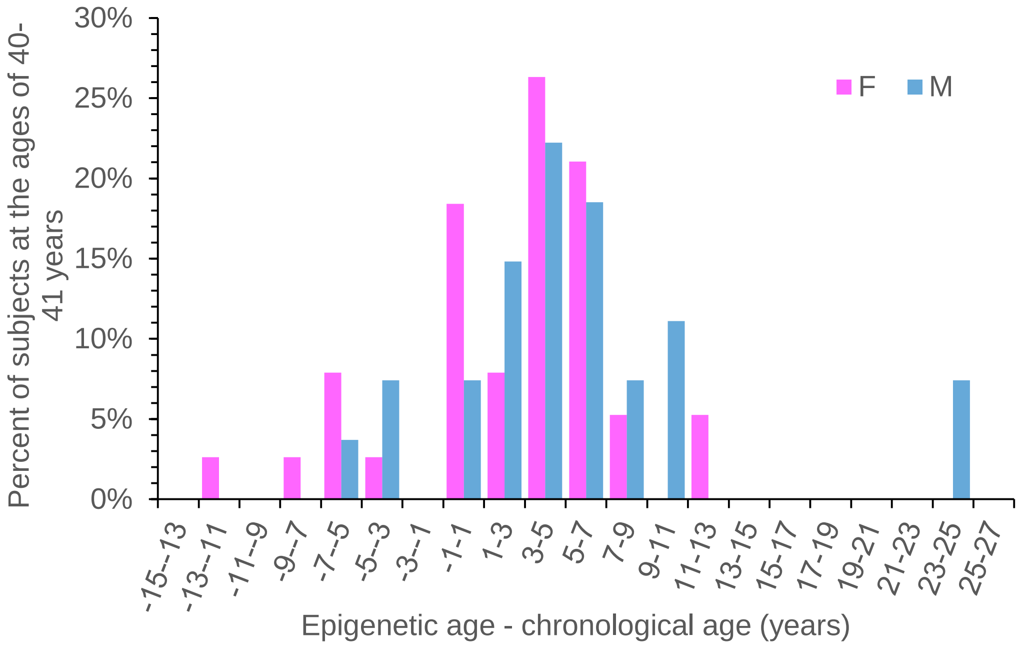

*Figure S1: Histogram of the difference between the calculated epigenetic age (by the 353 CpG sites of Horvath's clock) and the chronological age of men (blue bars) and women (purple bars) at the age of 40-41 years.*

#### 2.2. Table S2 and S3: Gender specific CpG sites with the highest inter-personal methylation variability

The CpG sites responssible for the epigenetic age variability within the healthy population were first examined in men and women seperatly. The 1441 healthy samples were divided according to gender. Each gender was divided to 8 chronological data sets by age groups spanning from the age of 40 to the age of 80 years**:** (40-45

(189); 46-50 (215); 51-55 (217); 56-60 (223); 61-65 (220); 66-70 (177); 71-75 (120); 76-80 (80), years (number of subjects)). At each chronological data set, we recorded the 20 CpG sites from Horvath's clock, with the highest inter-individual variability (standard deviation, SD, in years). We then identified the CpG sites out of the 20, which were also consistently present in at least 7 of the 8 chronological age data sets of the healthy cohort, for each gender. Ten out of the selected 20 CpG sites were found in at least 7 age groups of the men cohort and eleven were found in at least 7 age groups of the women cohort. These CpG sites and their associated genes are listed in the order of their SD values (highest to lowest) in table S2 (for men) and S3 (for women). The CpG sites that are shared by men and women are associated with the genes SCGN, NHLRC1, MIR7-3HG, FZD9, SCAP, REEP1, FXN, NDUFS5, B3GALT6 and CSNK1D. All had the same or almost the same SD value for both sexes. The only CpG site that entered the top 20 most variable CpG sites in 7 age groups in women but not in men, was the one associated with the SLC9A3R2 gene.

Table S2 (men, blue) and S3 (women, orange): Gender specific CpG sites and their related genes with the highest inter-personal methylation variability

| Men | | | | |
|---|---|---|---|---|
| **Gene Symbol** | **Product** | **CpGmarker** | **Contribution to epigenetic age (years)*** | **SD of the age contribution (years)** |
| SCGN | secretagogin precursor | cg06493994 | 7.6 | 2.1 |
| NHLRC1 | malin | cg22736354 | 9.3 | 1.8 |
| MIR7-3HG | hypothetical protein LOC284424 | cg02479575 | 1.7 | 1.0 |
| FZD9 | frizzled 9 | cg20692569 | 6.7 | 0.9 |
| SCAP | SREBP cleavage-activating protein | cg26614073 | -4.5 | 0.7 |
| REEP1 | receptor expression enhancing protein 1 | cg01968178 | 2.6 | 0.7 |
| FXN | frataxin isoform 1 preproprotein | cg07158339 | -4.5 | 0.7 |
| NDUFS5 | NADH dehydrogenase (ubiquinone) Fe-S protein 5 | cg07388493 | -4.6 | 0.7 |
| B3GALT6 | betaGal beta 1;3-galactosyltransferase polypeptide 6 | cg19945840 | 5.6 | 0.7 |
| CSNK1D | casein kinase 1; delta isoform 1 | cg19761273 | -4.0 | 0.7 |
| **Women** | | | | |
| **Gene Symbol** | **Product** | **CpGmarker** | **Contribution to epigenetic age (years)*** | **Women SD of the age contri (years)** |
| SCGN | secretagogin precursor | cg06493994 | 6.8 | 2.0 |
| NHLRC1 | malin | cg22736354 | 9.2 | 1.7 |
| MIR7-3HG | hypothetical protein LOC284424 | cg02479575 | 1.9 | 1.0 |
| SLC9A3R2 | solute carrier family 9 isoform 3 regulator 2 | cg08331960 | -3.6 | 0.9 |
| FZD9 | frizzled 9 | cg20692569 | 6.9 | 0.9 |
| B3GALT6 | betaGal beta 1;3-galactosyltransferase polypeptide 6 | cg19945840 | 5.7 | 0.7 |
| SCAP | SREBP cleavage-activating protein | cg26614073 | -4.8 | 0.7 |
| REEP1 | receptor expression enhancing protein 1 | cg01968178 | 2.4 | 0.7 |
| FXN | frataxin isoform 1 preproprotein | cg07158339 | -4.6 | 0.7 |
| CSNK1D | casein kinase 1; delta isoform 1 | cg19761273 | -4.5 | 0.7 |
| NDUFS5 | NADH dehydrogenase (ubiquinone) Fe-S protein 5 | cg07388493 | -4.9 | 0.6 |

2.3. Gender differences in the β values of the nine selected CpG sites

We tested for gender differences of individuals from the same age group in the β values of the nine selected most variable CpG sites. Statistics was performed using Statistica 7.1 software. Variables were tested using the parametric MANOVA (multiple analysis of variance; Friedman test was used when data was significantly different from normal distribution). We revealed a significant gender effect in the β values within the same chronological age group (ANOVA with repeated measure, F1 52=25, $p<0.0001$).

2.4. Figure S2: Correlation of methylation level between each selected CpG site and proximate CpG sites

The Illumina methylation arrays contain several CpG sites that are related to a single gene. However, only one CpG site per gene is included as a component in the epigenetic age calculators. We therefore examined how the CpG sites included in the epigenetic age calculation correlate with other CpG sites of the same gene. As shown in figure S2, the CpG site cg06493994 (yellow), related to the secretagogin gene, which is included in the epigenetic age clock, is highly correlated to other CpG sites with close proximity (red box), on the same gene. This indicates that the methylation level of the site included in the calculation of the epigenetic age provides good representation of the methylation status of neighboring sites and is therefore likely to be related to gene expression, if it resides on the promoter or another regulatory region. Other examples for the correlation between the methylation of a CpG site with the most inter-personal variability, which is included in the age calculator and other, neighboring, CpG sites, that are related to the same gene, show similar correlation pattern (supplementary file 5).

**A**

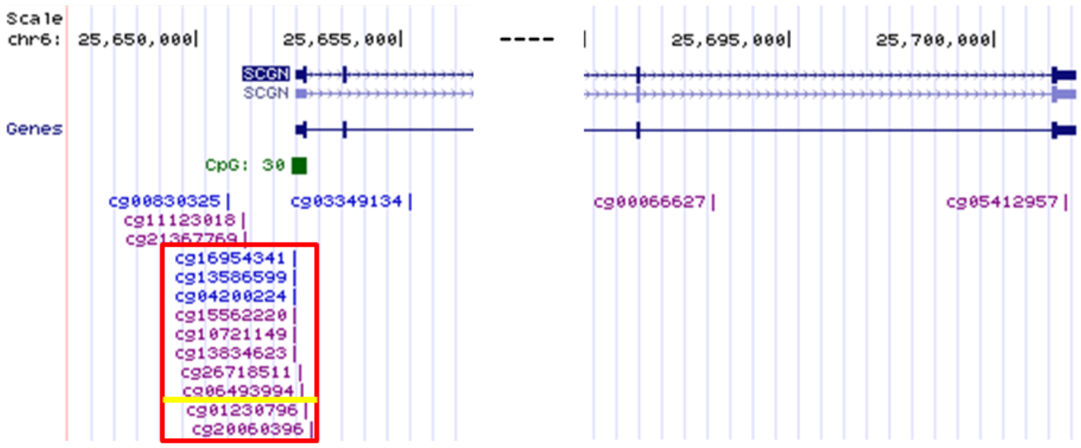

| **B** | cg00066627 | cg11123018 | cg05412957 | cg21367769 | cg03349134 | cg20060396 | cg13586599 | cg01230796 | cg06493994 | cg26718511 | cg15562220 | cg16954341 | cg10721149 | cg04200224 | cg13834623 |
|---|---|---|---|---|---|---|---|---|---|---|---|---|---|---|---|
| cg00066627 | 1.00 | 0.98 | 0.58 | 0.72 | -0.28 | -0.95 | -0.72 | -0.91 | -0.93 | -0.92 | -0.84 | -0.82 | -0.84 | -0.87 | -0.89 |
| cg11123018 | 0.98 | 1.00 | 0.57 | 0.64 | -0.37 | -0.97 | -0.79 | -0.96 | -0.97 | -0.96 | -0.88 | -0.88 | -0.90 | -0.92 | -0.93 |
| cg05412957 | 0.58 | 0.57 | 1.00 | 0.46 | -0.31 | -0.60 | -0.36 | -0.50 | -0.51 | -0.53 | -0.43 | -0.45 | -0.45 | -0.48 | -0.49 |
| cg21367769 | 0.72 | 0.64 | 0.46 | 1.00 | 0.00 | -0.63 | -0.22 | -0.45 | -0.49 | -0.48 | -0.41 | -0.33 | -0.38 | -0.40 | -0.43 |
| cg03349134 | -0.28 | -0.37 | -0.31 | 0.00 | 1.00 | 0.38 | 0.20 | 0.39 | 0.38 | 0.38 | 0.17 | 0.32 | 0.26 | 0.29 | 0.27 |
| cg20060396 | -0.95 | -0.97 | -0.60 | -0.63 | 0.38 | 1.00 | 0.79 | 0.94 | 0.94 | 0.94 | 0.86 | 0.88 | 0.87 | 0.89 | 0.91 |
| cg13586599 | -0.72 | -0.79 | -0.36 | -0.22 | 0.20 | 0.79 | 1.00 | 0.91 | 0.90 | 0.90 | 0.96 | 0.97 | 0.95 | 0.95 | 0.94 |
| cg01230796 | -0.91 | -0.96 | -0.50 | -0.45 | 0.39 | 0.94 | 0.91 | 1.00 | 0.99 | 0.99 | 0.95 | 0.97 | 0.97 | 0.98 | 0.98 |
| cg06493994 | -0.93 | -0.97 | -0.51 | -0.49 | 0.38 | 0.94 | 0.90 | 0.99 | 1.00 | 0.99 | 0.94 | 0.96 | 0.96 | 0.97 | 0.98 |
| cg26718511 | -0.92 | -0.96 | -0.53 | -0.48 | 0.38 | 0.94 | 0.90 | 0.99 | 0.99 | 1.00 | 0.95 | 0.96 | 0.96 | 0.98 | 0.98 |
| cg15562220 | -0.84 | -0.88 | -0.43 | -0.41 | 0.17 | 0.86 | 0.96 | 0.95 | 0.94 | 0.95 | 1.00 | 0.98 | 0.98 | 0.98 | 0.98 |
| cg16954341 | -0.82 | -0.88 | -0.45 | -0.33 | 0.32 | 0.88 | 0.97 | 0.97 | 0.96 | 0.96 | 0.98 | 1.00 | 0.98 | 0.98 | 0.98 |
| cg10721149 | -0.84 | -0.90 | -0.45 | -0.38 | 0.26 | 0.87 | 0.95 | 0.97 | 0.96 | 0.96 | 0.98 | 0.98 | 1.00 | 0.99 | 0.99 |
| cg04200224 | -0.87 | -0.92 | -0.48 | -0.40 | 0.29 | 0.89 | 0.95 | 0.98 | 0.97 | 0.98 | 0.98 | 0.98 | 0.99 | 1.00 | 0.99 |
| cg13834623 | -0.89 | -0.93 | -0.49 | -0.43 | 0.27 | 0.91 | 0.94 | 0.98 | 0.98 | 0.98 | 0.98 | 0.98 | 0.99 | 0.99 | 1.00 |

*Figure S2: Correlations of key CpG sites: (A) a map of two fragments from the SCGN gene from the NCBI wesite; blue cg are for non- methylated site, purple cg are for partly methylated site in GM12878 B-lymphocyte (B) Correlation matrix between the probe (CpG site) included in Horvath's clock (yellow line), which is related to the secretagogine (SCGN) gene and other CpG sites on the same gene, which are presented in Illumina's HumanMethylation450 BeadChip array. The red box includes a group of CpG sites with a high correlation to the CpG site from the aging algorithm (yellow line).*

### 2.5. Table S4: The dependency of population size on the selection of the most interpersonal variable CpG sites by bootstrapping simulation

The dependency of the selection of the most interpersonal variable CpG sites, on the size of the healthy population, was examined by statistical analysis, based on bootstrapping [6], [7] simulations (×1000), which were carried out with random sampling of 1441 subjects (equal to the number of subjects in the cohort). The same 9 most variable CpG probes were reselected. Eight out of the nine probes were reselected in more than 80% of the simulations, indicating a confidence of 80% or higher. Secretagogin was reselected at 997 out of 1000 simulations (99.7%), malin at 97.6%. SREBP cleavage-activating protein, NADH dehydrogenase (ubiquinone) Fe-S protein 5, frataxin, were all selected in more than 90% of the simulations; frizzled 9 and MIR-7 in more than 80% and the 9$^{th}$ probe (receptor expression enhancing protein 1) was found in 67.7 % of the simulations (table S4).

Table S4, The dependency of population size on the selection of the most interpersonally variable CpG sites by bootstrapping simulation

| Related gene symbol | Related Gene definition/ product | Illumina's CpG ID | % Simulations, in which the specific CpG site was selected as the most variable |
|---|---|---|---|
| SCGN | Secretagogin | cg06493994 | 99.7 |
| NHLRC1 | Malin | cg22736354 | 97.6 |
| MIR7-3HG | MIR7-3 Host Gene | cg02479575 | 81 |
| FZD9 | Frizzled 9 | cg20692569 | 88.3 |
| SCAP | SREBP cleavage-activating protein | cg26614073 | 97.2 |
| REEP1 | Receptor expression enhancing protein 1 | cg01968178 | 67.7 |
| CSNK1D | Casein kinase 1 delta | cg19761273 | 80.3 |
| FXN | Frataxin | cg07158339 | 90.3 |
| NDUFS5 | NADH dehydrogenase Fe-S protein 5 | cg07388493 | 92.2 |

### 2.6. Table S5: Bootstrapping simulation for the dependency of the CpG sites chosen by the greedy algorism on the population size

In the main text we have used a greedy algorithm in order to find out which components (CpG sites) are most responsible for age acceleration in the healthy cohort (figure 3). In the process of such analysis the epigenetic age of individuals in the group of ‘epigenetically old’ subjects are recalculated after setting the methylation level of chosen CpG sites, to their average value. The first CpG site chosen, is the one responsible for the highest number of individuals shifting from the “epigenetically old” epigenetic age group to the average + 1SD epigenetic age group (figure 1) after setting its value to the average contribution (in years). The second CpG site chosen is the one responsible for the second highest number of individuals shifting from the “epigenetically old” group to the average + 1SD group following normalization of both consecutive CpG sites, and so on.

The dependence of the results from the greedy algorithm on the population size was statistically analysed by bootstrapping (×1000) [6], [7] of 135 subjects residing in the “epigenetically old" healthy cohort. The key agers of the healthy population, secretagogin and malin were found in 100% of the simulations; MIR-7 and SREBP cleavage-activating protein in more than 95% and NADH dehydrogenase (ubiquinone) Fe-S protein 5, frataxin and receptor expression enhancing protein 1 in more than 55% of the simulations (table S9). These results emphasize how robust these CpG sites are as age accelerators and that the result is independent of the sample size.

Table S5: Statistical simulation for the dependency of the selected CpG sites as age accelerators, on the population size

| Related gene symbol | Related Gene definition/ product | Illumina's CpG ID | % Simulations, in which the specific CpG site was selected as accelerating/ decelerating |
|---|---|---|---|
| SCGN | Secretagogin | cg06493994 | 100 |
| NHLRC1 | Malin | cg22736354 | 100 |
| MIR7-3HG | MIR7-3 Host Gene | cg02479575 | 100 |
| SCAP | SREBP cleavage-activating protein | cg26614073 | 95.3 |
| REEP1 | Receptor expression enhancing protein 1 | cg01968178 | 55.7 |
| CSNK1D | Casein kinase 1 delta | cg19761273 | 13.3 |
| FXN | Frataxin | cg07158339 | 62.6 |
| NDUFS5 | NADH dehydrogenase Fe-S protein 5 | cg07388493 | 68.1 |

2.7. Tables S6, S7 and S8: CpG sites with the highest inter-personal methylation variability in the diabetic and in the healthy cohorts

The interpersonal variability in the diabetic cohort was examined by looking for the CpG sites that were the most variable among subjects in all chronological age groups, in terms of years added/subtracted by that site to/from the total epigenetic age. Tables S6-S8 show the similarity in the list of genes related to the most variable CpG sites across the diabetic (T1D and T2D) and healthy cohorts, with some difference in the order of the size of the standard deviation (in years). Secretagogin and malin were found to be the genes related to the leading most variable CpG sites in all of the four different populations: T1D alone, T2D alone, T1D united with T2D and healthy (tables S6-S8, indicated in green). Whereas, PRKC Apoptosis WT1 Regulator (PAWR) and L-pipecolic acid oxidase (PIPOX) are genes related to the top 5 most variable CpG sites that are common only to T1D and T2D (tables S6-S8, indicated in orange). Six and four CpG sites out of the nine most variable sites of the healthy population are shared by T2D and T1D, respectively (tables S6- S8, indicated in green and blue).

Table S6: CpG sites with the highest inter-personal methylation variability in the type 2 diabetes mellitus cohort

| T2D | | | | |
|---|---|---|---|---|
| **Related gene symbol** | **Related Gene definition/ product** | **Illumina's CpG ID** | **Contribution to epigenetic age (years)*** | **SD of the age contribution (years)** |
| SCGN | Secretagogin | cg06493994 | 9.3 | 1.3 |
| NHLRC1 | Malin | cg22736354 | 11.7 | 1.2 |
| DPP8 | dipeptidyl peptidase 8 isoform 3 | cg06993413 | 2.7 | 1.2 |
| PAWR | PRKC Apoptosis WT1 Regulator | cg00864867 | 1.7 | 0.7 |
| CSNK1D | Casein kinase 1 delta | cg19761273 | -4.2 | 0.6 |
| PIPOX | L-pipecolic acid oxidase | cg06144905 | 3.9 | 0.6 |
| NDUFS5 | NADH dehydrogenase (ubiquinone) Fe-S protein 5 | cg07388493 | -4.8 | 0.6 |
| SCAP | SREBP cleavage-activating protein | cg26614073 | -4.6 | 0.6 |
| FZD9 | Frizzled 9 | cg20692569 | 7.2 | 0.6 |

**genes related to CpG sites that are common to all four populations: T1D alone, T2D alone, T1D and T2D united and healthy, are indicated in green, genes that are common only to T1D and T2D but not to the healthy population are indicated in orange, genes that are common to T1D or T2D and to the healthy population are indicated in blue*

Table S7: CpG sites with the highest inter-personal methylation variability in T1D cohort

| T1D | | | | |
|---|---|---|---|---|
| **Related gene symbol** | **Related Gene definition/ product** | **Illumina's CpG ID** | **Contribution to epigenetic age (years)*** | **SD of the age contribution (years)** |
| NHLRC1 | Malin | cg22736354 | 8.734 | 1.012 |
| PAWR | PRKC Apoptosis WT1 Regulator | cg00864867 | 3.785 | 0.700 |
| SCGN | Secretagogin | cg06493994 | 6.882 | 0.642 |
| FZD9 | Frizzled 9 | cg20692569 | 6.389 | 0.534 |
| PIPOX | L-pipecolic acid oxidase | cg06144905 | 3.051 | 0.514 |
| FXN | Frataxin | cg07158339 | -5.547 | 0.503 |
| SSRP1 | structure specific recognition protein 1 | cg01511567 | -2.879 | 0.480 |
| FLJ21839 | hypothetical protein LOC60509 | cg14424579 | 2.290 | 0.440 |

**genes related to CpG sites that are common to all four populations: T1D alone, T2D alone, T1D and T2D united and healthy, are indicated in green, genes that are common only to T1D and T2D but not to the healthy population are indicated in orange, genes that are common to T1D or T2D and to the healthy population are indicated in blue*

Table S8: A comparison between CpG sites with the highest inter-personal methylation variability in T1D, T1D, T1D+T2D and healthy cohorts

| T1D | T2D | T1D+T2D | Healthy |
|---|---|---|---|
| NHLRC1 | SCGN | NHLRC1 | SCGN |
| PAWR | NHLRC1 | SCGN | NHLRC1 |
| SCGN | DPP8 | PAWR | MIR7-3HG |
| FZD9 | PAWR | PIPOX | FZD9 |
| PIPOX | CSNK1D | FZD9 | SCAP |
| FXN | PIPOX | | REEP1 |
| SSRP1 | NDUFS5 | | CSNK1D |
| FLJ21839 | SCAP | | FXN |
| | FZD9 | | NDUFS5 |

**genes related to CpG sites that are common to all four populations: T1D alone, T2D alone, T1D and T2D united and healthy, are indicated in green, genes that are common only to T1D and T2D but not to the healthy population are indicated in orange, genes that are common to T1D or T2D and to the healthy population are indicated in blue*

References:

[1] S. Horvath, "DNA methylation age of human tissues and cell types," *Genome*

*Biol.*, vol. 14, no. 10, 2013, doi: 10.1186/gb-2013-14-10-r115.
